# What We Miss on the Road: Visual Attention and Species Detectability in Roadkill Surveys

**DOI:** 10.1101/2025.07.15.664863

**Authors:** Annaëlle Bénard, Thierry Lengagne

## Abstract

Wildlife-vehicle collisions pose a substantial threat to biodiversity, and yet roadkill counts—one the main ways used to quantify mortality—are often susceptible to underestimate true collision rates due to imperfect detection. This study investigates how carcass size and survey methodology affect detection rates during roadkill surveys. Using taxidermied specimens placed along roads, we compare roadkill detection in standardized surveys vs. opportunistic conditions that mimic citizen science data collection, in which observers are not asked to focus on roadkill search alongside roads prior to the survey. Results show that detection probability increases with body mass but declines sharply when observers are not explicitly focused on locating roadkill, with informed participants up to 27 times more likely to detect carcasses than uninformed ones. Even large-bodied species, such as red foxes and European badgers, were frequently missed in opportunistic contexts. By applying species-specific detection probabilities and carcass persistence estimates to a regional citizen science roadkill dataset (Faune-AuRA), we reveal that reported carcass counts represent as little as 0.8–5% of estimated collision numbers. These results demonstrate that roadkill counts, regardless of survey methodology, may substantially underestimate road mortality and distort assessments of species-specific vulnerability in transportation ecology.

## 1. Introduction

Terrestrial transportation contributes to the on-going global biodiversity decline by threatening the fitness of individuals (Forman & Alexander, 1998). In particular, building roads result in habitat loss and fragmentation that disrupt normal ecological processes (Seiler, 2001), and the acoustic, chemical and light pollution generated by road traffic are detrimental to the survival of many species (Bissonette & Rosa, 2009; Eigenbrod et al., 2009; Forman, 2000). Collisions between wildlife and vehicles are a direct source of mortality that potentially affects all terrestrial species (Olson et al., 2014; Swinnen et al., 2022) and may threaten population viability in many cases (Grilo et al., 2021; Moore et al., 2023).

Collisions occur at uneven rates depending on the species, location, season and time of day (Hastings et al., 2019; Ignatavičius et al., 2020; Quiles, 2018; Teixeira et al., 2013). The impact of road mortality could be as high as 194 millions of mammals and 29 millions of birds annually in Europe (Grilo et al., 2020). There is an urgent need to quantify this mortality across species and develop effective mitigation strategies. While fences and structures such as underpasses or overpasses enable safe crossings and help wildlife avoid collisions with vehicles (Bond et al., 2008), such measures come at a high cost. There is therefore a need to deepen our understanding of road mortality’s impact on wildlife populations—and especially for emblematic species that are likely to draw public and stakeholder attention, and thus drive momentum for action.

Collecting wildlife-collision data is essential towards this goal. It is often gathered through standardized surveys that count roadkill carcasses along designated routes (e.g., Canova & Balestrieri, 2019). In recent years, large-scale citizen science initiatives have emerged as an alternative where contributors opportunistically report carcass sightings on roadways. These projects have grown in popularity, with some reaching thousands of contributors and generating over ten of thousands of reports for a wide range of species, and at a fraction of the cost of professional surveys (Swinnen et al., 2022). However, carcass counts are prone to systematic underestimation: roadkill remains on roads only briefly, and detection is usually imperfect. Reports submitted by volunteers also introduce additional sources of bias, including misidentification of species and unmeasured, uneven sampling effort. Despite these limitations, multiple authors have argued that citizen science roadkill data can provide valuable insights into collision rates (Petrovan et al., 2020; Shin et al., 2022; Waetjen & Shilling, 2017).

In recent years, increasing attention has been given to identifying biases in standardized roadkill surveys. For instance, studies have estimated carcasses persistence time on roads following a collision (S. M. Santos et al., 2011), which in turn allows the selection of appropriate monitoring frequencies to minimize bias (Henry et al., 2021). Detection probabilities have also been assessed, often by comparing the number of carcasses observed from a vehicle to those detected by observers on foot (Table 1). Survey recommendations tailored to the species and context—such as mode of travel (on foot, by bicycle, or by car) and travel speed—can help reduce the number of missed carcasses during professional monitoring schemes (Collinson et al., 2014). However, while some estimates of detection rates exists in the literature across broad categories of body mass or taxa (Gerow et al., 2010; R. A. L. Santos et al., 2016; Teixeira et al., 2013), few studies have examined species-level detection patterns in detail. As a result, there remain limited resources to accurately adjust carcass counts in a planned survey by accounting for detection probabilities for a given species.

**Table 1.** Review of the published roadkill detection probabilities estimated during standardized road surveys. The detectability of roadkill from a moving vehicle is more often estimated by either using decoys placed on the road or by comparing the results of surveys performed from a vehicle to surveys performed on foot, assuming that foot survey detect 100% of the roadkill present on the road. In addition to size and taxa, the detectability of roadkill varies with the driving speed of the survey vehicle and the level of experience of the observer (Collinson et al., 2014).

| Authors | P<sub>detection</sub> (95% CI) | Method |
| --- | --- | --- |
| Collinson et al. (2014) | 88.8% (83.8-92.8) | Using painted rubber to simulate flattened roadkill placed on 1-km road transects |
| Gerow et al. (2010) | Amphibians: 4.8% (4-5.6)<br>Reptiles: 3.7% (2.6-4.8)<br>Birds: 34.5% (28.3-40.7)<br>Mammals: 10.3% (7.6-13) | Comparing the number of carcasses recorded during a driving survey to the number detected during a concurrent walking survey |
| Santos et al. (2016) | 7% (2-15) for animals <100g<br>13.3% (4-29) for animals >100g |  |
| Teixeira et al. (2013) | Reptiles: 20% (7-33)<br>Birds: 5% (4.3-5.7)<br>Mammals: 47% (34-60) |  |

In addition, little attention has been given to the specific case of opportunistic citizen science datasets. While carcass persistence time has no reason to differ depending on survey type, there are multiple reasons to believe that the roadkill detectability estimates during surveys may not be easily transferable from standardized roadkill surveys to opportunistic data. In citizen science projects, observers typically drive without a strong focus on detecting roadkill. This difference in attentional engagement can be re-framed using psychological research on attention: the brain is only capable of processing a limited amount of information at any given time, meaning that attention is a constrained resource. Only a fraction of the visual input captured by the eyes is actually analyzed by the brain (Dukas, 2004). In visual search tasks—such as locating carcasses on roadways—being explicitly instructed to focus on the objects being searched can substantially increase detection success, by engaging in top-down attentional control (Chen & Zelinsky, 2006). Moreover, goal-directed search, such as during standardized surveys, may enhance sensitivity to low-salience features: observers in standardized surveys may not only detect more carcasses overall—because they focus on the object of their search—they may also be proportionally better at detecting smaller or more cryptic species. In other words, survey context (opportunistic vs. standardized survey) may not only lead to differences in detection rates for a given species, but it may also modulate the very relationship between carcass characteristics and detection rates (Wolfe & Horowitz, 2017).

In this study, we investigate roadkill detection rates with a particular focus on animal size and features, as well as the observer’s attentional state. Our primary objective is to develop a predictive model of detection probability using roadkill body mass as a continuous variable and other species-specific features, in contrast to previous studies that typically rely on broad categorical groupings. Additionally, we examine the transferability of detection rates between standardized roadkill surveys—where observers are explicitly instructed to search for carcasses and remain attentive during the whole drive—and opportunistic datasets in which observers report roadkill without focusing their attention on the visual search. We hypothesize that (1) detection rates are substantially lower in opportunistic data collection, compared to standardized surveys, and (2) visual search in standardized surveys is not only more efficient but also qualitatively different, potentially yielding proportionally higher detection rates for smaller or cryptic species. Finally, we apply our estimates of detection rates to a citizen science dataset consisting of opportunistic reports of roadkill across the Auvergne-Rhône-Alpes region in France. By assuming a fixed survey rate, adjusting for under-reporting due to limited carcass persistence (Benard, 2023) and imperfect detection, as estimated in this study, we demonstrate how little opportunistic reports of carcass presence on roads may represent true carcass counts. At the same time, these estimates also aim at highlighting the potential for opportunistic data collection schemes to generate broad estimates of collision rates across a wide range of species.

## 2. Material and Methods

### 2.1 Roadkill decoys

We used taxidermied animals to simulate roadkill during field experiments. This method ensured that each specimen closely matched the size and coloration of fresh roadkill, while enabling consistent comparisons between informed (as under standardized survey conditions) and uninformed (as during opportunistic data collection) detection rates by using identical animal models. We obtained 25 taxidermied individuals of 11 different species of birds and mammals, sourced either from private collectors or through professional preservation of carcasses donated by a wildlife rehabilitation center (see Appendix A1 for full species list).

During the trials, specimens were placed in randomized order along the road shoulders or at the boundary between the road surface and shoulder, to ensure visibility and avoid obstruction by vegetation. Each was anchored using camping pegs. For safety reasons, no specimens were placed directly on the roadway. Some specimens were lost or removed during the experiments (including two snakes and ten small passerines) and were excluded from subsequent analyses, leaving 20 specimens in total.

### 2.2 Roadkill detectability experiments

#### Informed search

We conducted a field experiment on the road network of the Ain department in central-eastern France. A 28 km circuit was defined, covering roads with traffic volumes ranging from under 500 to over 9989 vehicles per day. Because the impact of observer speed on detection rates were not the focus of the present study, all specimens were placed along sections of road with posted speed limits of 80 km.h^-1^, except for one red squirrel positioned on a 90 km.h^-1^ stretch.

Twenty specimens were placed on the right-hand side of the road (the driving side for participants, see fig. A2) and spaced at least 700 meters apart. To deter theft and monitor potential removal, the largest specimens were equipped with camera traps. The experiment took place on an overcast morning with moderate rainfall (2 to 8 mm.h^-1^) and no fog. We recruited N = 17 volunteer drivers with no prior experience in roadkill surveys, primarily through the Bird Protection League (LPO) and other student naturalist associations. Participants therefore had a likely above-average familiarity with common wildlife species, which facilitated the analysis of the data they produced as they usually identified the species correctly even from a moving vehicle.

Each participant was informed of the task to be performed, and asked to drove their personal vehicle along the route at or near the posted speed limit. Their departure and arrival times were recorded to ensure compliance, with at least a 10-minute interval between successive drivers. Participants were asked to report detected species in two stages: once at a mid-point stop (after 14 km and passing 10 specimens), and again at the end of the circuit. These lists were compiled under the supervision of a field assistant to minimize ambiguous reports.

#### Naive (uninformed) search

To evaluate roadkill detection rates among drivers not actively searching for carcasses, we conducted two experimental sessions during key events organized by the Bird Protection League (LPO): the general assembly (June 2022; N = 62 attendees) and the board of directors meeting (October 2022; N = 17 attendees). These sessions were selected because LPO members represent an overwhelming majority of the contributors to the Faune-AuRA citizen-science roadkill program (Faune France, 2018), thus aligning with our goal of assessing typical observer performance within that framework. Specimens were placed along the final 5 km of road leading to each event’s venue (posted speed limits: 80 km.h^-1^), covering all possible access routes and ensuring consistent species exposure across itineraries (Fig. A3).

Specimen placement occurred at least 30 minutes before the arrival of the first attendees. In the first session, specimens included red squirrels (*Sciurus vulgaris*) and red foxes (*Vulpes vulpes*). In the second, we added diurnal raptors of similar size—the European honey buzzard (*Pernis apivorus*) and common buzzard (*Buteo buteo*)—as well as pigeons (*Columba livia*). Upon arrival, participants were invited to anonymously fill out forms in which they reported whether or not they had seen roadkill in the final 5 km of their journey, and indicating their seat in the vehicle (driver or passenger). Survey options listed all deployed species as well as a control species (badger) that was not present and used to identify potential false positives (Bhattacherjee, 2012).

### 2.3 Data analysis

### Roadkill detection rates

Species’ mean body masses were obtained from publicly available databases (Herberstein et al., 2022; Tobias et al., 2022; Tranquillo et al., 2024, MammalBase, 2024). To understand the relationship between specimen detectability, body size, taxonomic group, and observer attentiveness, we constructed a mixed-effects logistic regression model using individual detection scores (R package *lme4*, Bates et al., 2015). Predictor variables included species’ mean body mass (continuous), observer status (informed or naive), taxonomic group (bird or mammal), and the observer’s seat in the vehicle (driver or passenger; all informed participants were drivers).

We evaluated body mass both as a linear and log-transformed predictor, retaining the best-fitting model based on the Akaike Information Criterion (AIC). Taxonomic group was included to control for variation in body mass and structural traits influencing detectability—mostly because birds often exhibit lower mass relative to volume compared to mammals. To account for the possibility that structural traits (such as feathers or fur) shape the size-detectability relationship, we included an interaction between taxonomic group and body mass. Additionally, to test whether goal-directed search alters the relation between body mass and detection efficiency in drivers, we included an interaction term between body mass and observer status. Observer identity was modeled as a random effect to account for individual-level variation. Model diagnostics were conducted using the R *DHARMa* package (Hartig, 2022), which provides validation tools for generalized mixed models.

#### Estimating the number of animal-vehicle collisions using citizen-science data

We extracted the roadkill reports from Faune-AuRA (a citizen science dataset of opportunistic reports of roadkill presence on roads in the AuRA region) for 4 species used in the *naive* experiments: red fox (*Vulpes vulpes*), red squirrel (*Sciurus vulgaris*), common wood pigeons (*Columba palumbus*) and common buzzard (*Buteo buteo*), for the year 2022. Following Teixeira *et al*. (2013), the roadkill rate *λ* (collision.day^-1^) can be estimated as 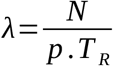, where N is the number of roadkill carcasses counted, p is the detection rate of the carcasses by the observer and T_R_ is a measure of the carcass persistence on the road defined as th*e* time needed to reduce the number of remaining carcasses on the road by 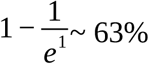. We used the predicted species-specific detection rates obtained in this study, alongside estimates of carcass persistence time in the region (T_R_) from Bénard et al. (2023), to estimate the total number of wildlife–vehicle collisions involving these species in the Auvergne-Rhône-Alpes road network in 2022. To estimate uncertainty in *λ*, we used a bootstrap procedure, resampling the datasets 1000 times and applying the equation at each iteration to construct empirical 95% confidence intervals. These calculations rely on key assumptions: (1) carcass persistence and observer detectability are the only sources of bias in collision reporting in Faune-AuRA and (2) all roads in the region are surveyed exactly once a day. These conditions are unlikely to be met in opportunistic citizen-science datasets, which usually exhibit variable observer effort, reporting biases in favor of certain species, and uneven spatial coverage in participants presence. Consequently, our estimates should be interpreted as conservative approximations of actual collision rates.

## 3. Results

Informed observers detected, on average, 73.8% of specimens (standard deviation: 12.8%). Detection scores dropped substantially in the uninformed experiments: for example, red foxes were detected 94% of the time by informed participants, but only 16.4% of the time by naive participants. None of the participants in the *naive* experiment reported having seen a badger (*i*.*e*., no false positives). The best-fitting logistic regression model included log-transformed body mass as a predictor (AIC_log_ = 459.9; AIC_linear_ = 462.3). Detection probability increased significantly with species body mass (β = 0.69, SE = 0.19, *p* < 0.001). At comparable weights, birds were more frequently detected than mammals (β = 0.98, SE = 0.32, *p* < 0.001), although the interaction between body mass and taxonomic group was not statistically significant (β = 0.39, SE = 0.32, *p* = 0.23). Informed observers were substantially more likely to detect specimens than uninformed ones (β = −3.29, SE = 0.41, *p* < 0.001), corresponding to an odds ratio of approximately 0.037. This implies that informed participants were about 27 times more likely to detect a given specimen than uninformed participants (Fig. 1). However, we find no significant interaction between body mass and observer status (β = −0.24, SE = 0.27, *p* = 0.38). Seat in the vehicle (driver vs. passenger) had no significant effect on detection probability either (β = −0.25, SE = 0.51, *p* = 0.62).

**Fig. 1.**
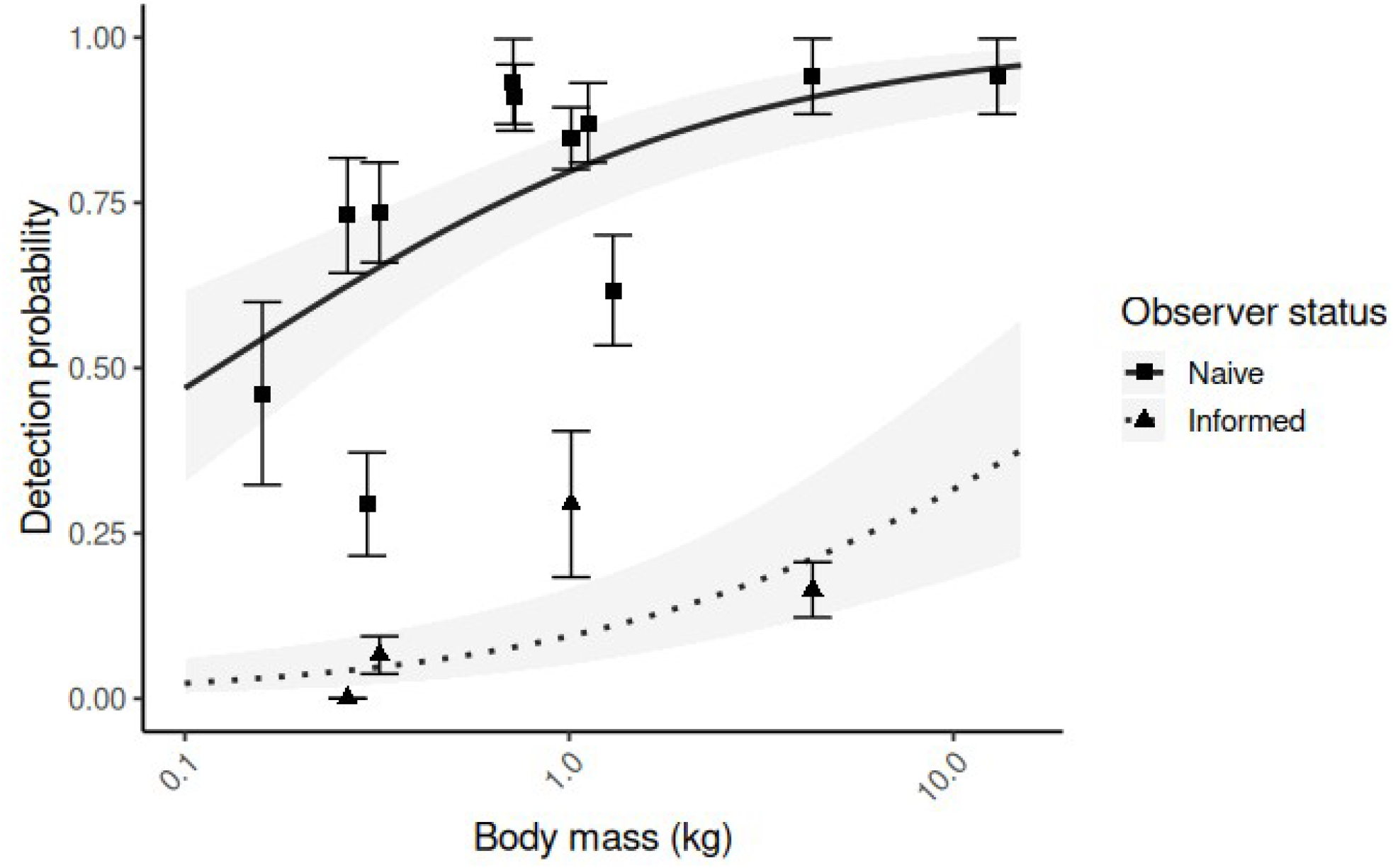
Predicted and observed roadkill detection probabilities for a driver. Points represent actual detection scores; lines and shaded areas show logistic regression predictions with 95% confidence intervals. Detection rates increase with species body mass but drop sharply when observers are not instructed to pay attention to roadkill prior to driving.

### Roadkill rate estimates from citizen-science data

We extracted a total of 607 reports of roadkill for *Vulpes vulpes*, 267 reports for *Buteo buteo*, 335 for *Sciurus vulgaris* and 63 for *Columba palumbus* from Faune-AuRA. Resulting collisions estimates are presented in table 3.

**Table 2.**
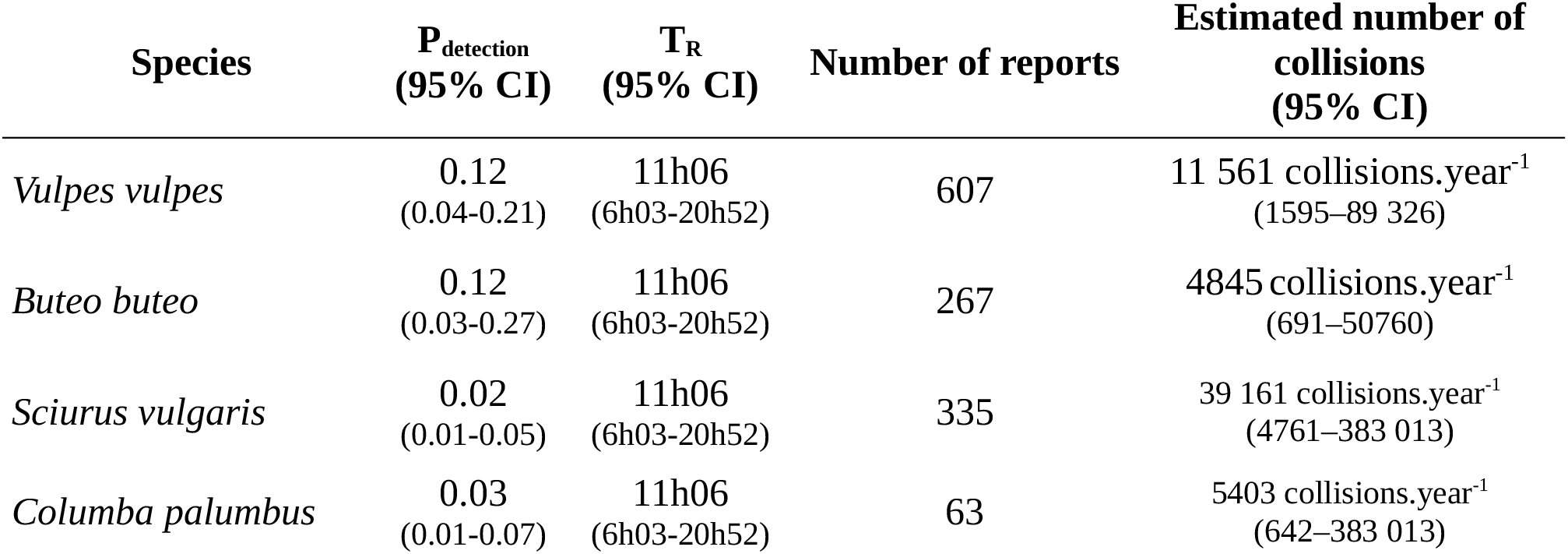
Estimated number of collisions with vehicles in the Auvergne-Rhône-Alpes region, France. We used citizen-science reports of roadkill from the Faune-AuRA database for 2022. Following Teixeira et al. (2013), we adjusted for the number of carcasses not reported with estimates of the detectability of animal carcasses in the context of citizen science (P_detection_) and of their persistence time on the road following a collision (T_R_, Bénard et al., 2023).

| Species | $P_{\text{detection}}$<br>(95% CI) | $T_R$<br>(95% CI) | Number of reports | Estimated number of<br>collisions<br>(95% CI) |
| --- | --- | --- | --- | --- |
| *Vulpes vulpes* | 0.12<br>(0.04-0.21) | 11h06<br>(6h03-20h52) | 607 | 11 561 collisions.year<sup>-1</sup><br>(1595–89 326) |
| *Buteo buteo* | 0.12<br>(0.03-0.27) | 11h06<br>(6h03-20h52) | 267 | 4845 collisions.year<sup>-1</sup><br>(691–50760) |
| *Sciurus vulgaris* | 0.02<br>(0.01-0.05) | 11h06<br>(6h03-20h52) | 335 | 39 161 collisions.year<sup>-1</sup><br>(4761–383 013) |
| *Columba palumbus* | 0.03<br>(0.01-0.07) | 11h06<br>(6h03-20h52) | 63 | 5403 collisions.year<sup>-1</sup><br>(642–383 013) |

## 4. Discussion

Imperfect detection is a central question in ecological monitoring. Any field survey aiming to assess species presence inevitably suffers from detection biases. To disentangle true presence from detection error, ecologists have developed new methods and modelling frameworks such as distance sampling, or occupancy models (Buckland et al., 2005; MacKenzie et al., 2002). In the context of roadkill surveys—typically classified as a form of visual encounter surveys—detection rates are particularly complex, as these surveys often span multiple specie and survey conditions simultaneously, and carcass condition may degrade over time further lowering detectability. Despite the critical role of roadkill count data in estimating mortality rates and identifying collision hotspots, methodological biases such as carcass persistence and imperfect detection have received comparatively little attention in the literature (R. A. L. Santos et al., 2016; Teixeira et al., 2013).

Detection probabilities reported in the literature vary widely across studies (see Table 1): visibility is at least influenced by carcass size, condition, and position on the road (Collinson et al., 2014). Standardized roadkill surveys are at risk of severely underestimating actual roadkill rate (R. A. L. Santos et al., 2016), and even trained observers may still miss carcasses—especially smaller ones and when surveying from a moving vehicle (Gerow et al., 2010). As a result, roadkill’s impact on wildlife populations may be perceived as less than it actually is (Teixeira et al., 2013).

Unsurprisingly, and consistent with previous findings, our results show that larger species are more easily detected than smaller ones. This also aligns with visual search literature, which identifies size as a key contributor to an object’s salience (Wolfe & Horowitz, 2017). We present here a more refined model for estimating detection rates in roadkill surveys by treating species’ average body mass as a continuous predictor—an advancement over previous categorical approaches. However, because our focus was on species-specific patterns, we omitted observer speed by simply ensuring all surveys were conducted on roads with a posted speed limit of 80 km.h^-1^. Therefore, while this study improves the consistency of available detection rates estimates, it also means that other influential variables—such as observer travel speed—remain unaccounted for and should be addressed in future work. In addition, although driver attentiveness has been extensively studied in the context of accident prevention (Larue et al., 2010; Lee, 2008; Underwood et al., 2002), little is known about the comparative detection performance of front-seat passengers versus drivers. Our data show no substantial difference in detection rates between the two groups; however, we were unable to assess passenger performance in informed search conditions. Given that standardized roadkill surveys are frequently conducted by passengers, to ensure safety, this gap in knowledge warrants further investigation.

By designing an experimental framework in which observers were not instructed to focus on roadkill detection prior to encountering simulated carcasses—that is, lacking top-down attentional control—we were able to estimate for the first the detection rates under opportunistic conditions. The effect of attentional engagement was striking: observers were up to 30 times less likely to detect carcasses when not actively searching. Even large-bodied species such as the European badger (∼10 kg) were frequently missed in these conditions: our findings show that imperfect detection remains widespread, even for conspicuous species, when observers are not actively looking or mentally prepared to identify roadkill. This challenges the notion that larger species may be reliably detected in citizen science road monitoring programs and has implications for other forms of opportunistic data collection as well, where the challenge of distinguishing true absences from false absences—often treated as pseudo-absences in modeling—has been widely discussed in recent years, particularly in relation to unmeasured sampling effort and imperfect detection (Dorazio, 2014; Kéry et al., 2010).

We find that the gap between the number of reports received under the opportunistic monitoring program and the estimates of actual road mortality is substantial. Smaller species, being less salient, exhibit even greater discrepancies between reported and actual collision rates than larger ones. For instance, we estimate that opportunistic reports of red foxes in the Faune-AuRA program represent only ∼5% of actual collisions. For red squirrels, this figure drops to ∼0.8%. These gaps are likely exacerbated by faster carcass disappearance for smaller species (S. M. Santos et al., 2011), which we did not model here. Reporting biases in citizen science programs may further skew species representation: participants may preferentially report charismatic or rare—those perceived as more “important” or interesting (August et al., 2020), introducing an additional layer of unpredictability in the relationship between reported and actual collision numbers. Consequently, not only do opportunistic roadkill reports likely mask an immense toll of transportation on wildlife population, they also distort the relative impact across species. The multiplication factors we calculated to scale from reports to estimated collisions are especially concerning, as some citizen science programs now exceed hundreds of thousands of reports (Swinnen et al., 2022; Faune-AuRA, 2023), suggesting that the true scale of roadkill-related losses may be vastly underestimated and largely overlooked by the ecological community.

## 5. Conclusions

Imperfect detection is a central concern in ecological monitoring, including roadkill surveys. Detection rates observed during standardized surveys vary widely across species and carcass sizes, highlighting how species traits influence visibility. Opportunistic datasets from citizen science programs suffer from similar detection issues. While attention in presence-only data is often directed toward spatial and temporal variation in sampling effort, the specific context in which opportunistic observations are collected—where drivers are not actively searching for roadkill—can dramatically reduce detection efficiency, even for medium-to large-bodied species. By accounting for this overlooked source of bias, it becomes clear that the proportion of citizen science reports relative to estimated collision numbers may conceal an alarming scale of wildlife mortality on road networks.

## 6. Ethical statement

The participants recruited in this experiment gave free and informed consent for their answers to be used for the purposes of this study. Surveys were entirely anonymous and no other identifying data was collected.

## 7. Acknowledgments

We thank the volunteers for participating in this study, as well as J. Barbe for helping in the experiments. We also thank the Tichodrome wildlife rehabilitation center for providing carcasses and J. Gilbert for his work in preserving the animals.

