## Appendix for "What We Miss on the Road: Visual Attention and Species Detectability in Roadkill Surveys"

A1: complete list of the decoy species – volume of the decoy. These taxidermies were used for the detectability experiments. Because the diurnal raptors (Red kite, Common buzzard, Honey buzzard, Marsh harrier) were very similar in coloration and volume, most participants could not accurately discriminate the species during the experiments. Consequently, they were pooled into a ‘Diurnal raptor’ category and given the same average body mass.

- Brown rat (*Rattus norvegicus) - 0.298 kg*
- Common buzzard *(Buteo buteo) – 1.012 kg*
- Common pheasant (*Phasianus colchicus) – 1.12 kg*
- Domestic pigeon *(Columba livia) – 0.265 kg*
- European badger (*Meles meles) – 13 kg*
- European hedgehog *(Erinaceus europaeus) - 0.723 kg*
- European honey buzzard (*Pernis apivorus) – 1.012 kg*
- Eurasian jay (*Garrulus glandarius) – 0.159 kg*
- European pine marten (*Martes martes) – 1.300 kg*
- Peregrine falcon (*Falco peregrinus) – 0.715 kg*
- Red fox *(Vulpes vulpes) – 4.300 kg*
- Red kite (*Milvus milvus) – 1.012 kg*
- Red squirrel (*Sciurus vulgaris) – 0.321 kg*
- Western marsh harrier *(Circus aeruginosus) – 1.012 kg*

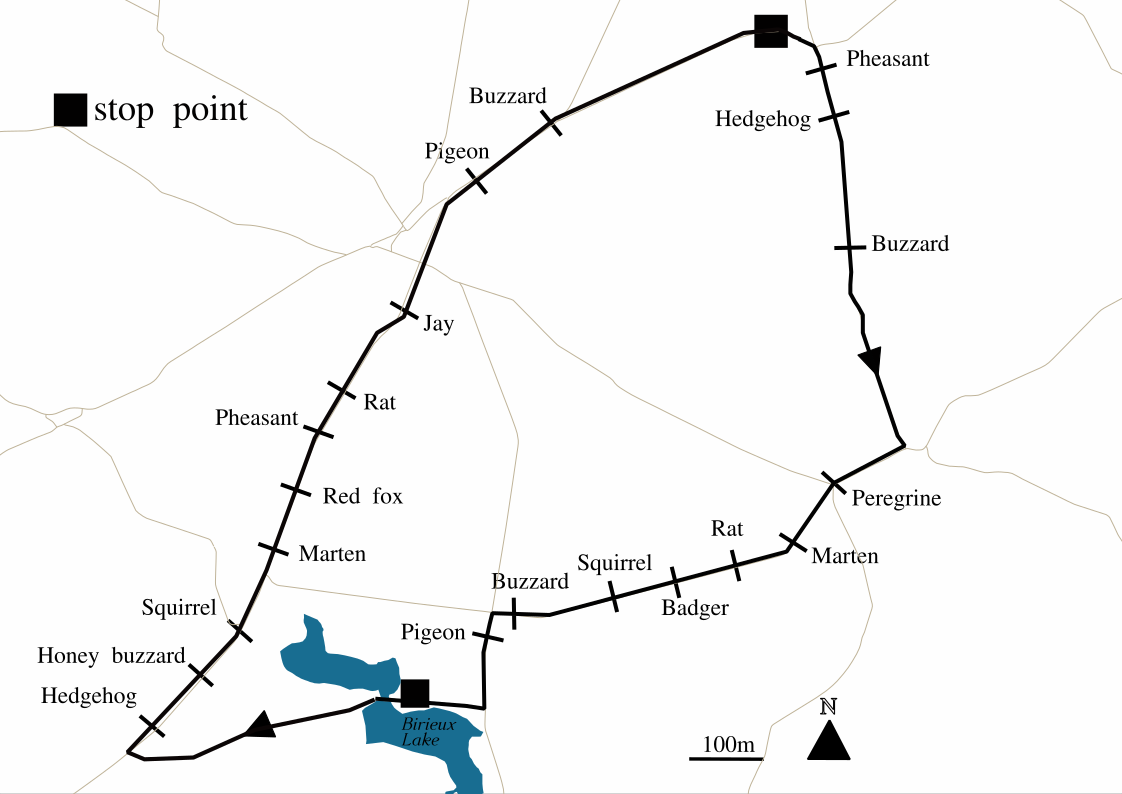
Figure A2: Map of the *informed experiment. To estimate the detection score of several species on the road, we placed the 20 taxidermies on a 28km circuit. We asked the participants to drive on the circuit with their vehicle and write down the list of species they had detected. Participants stopped twice during the experiment to write at determined stop points (■), such that they would have to remember a maximum of 10 species (if all were detected) before writing the list.*

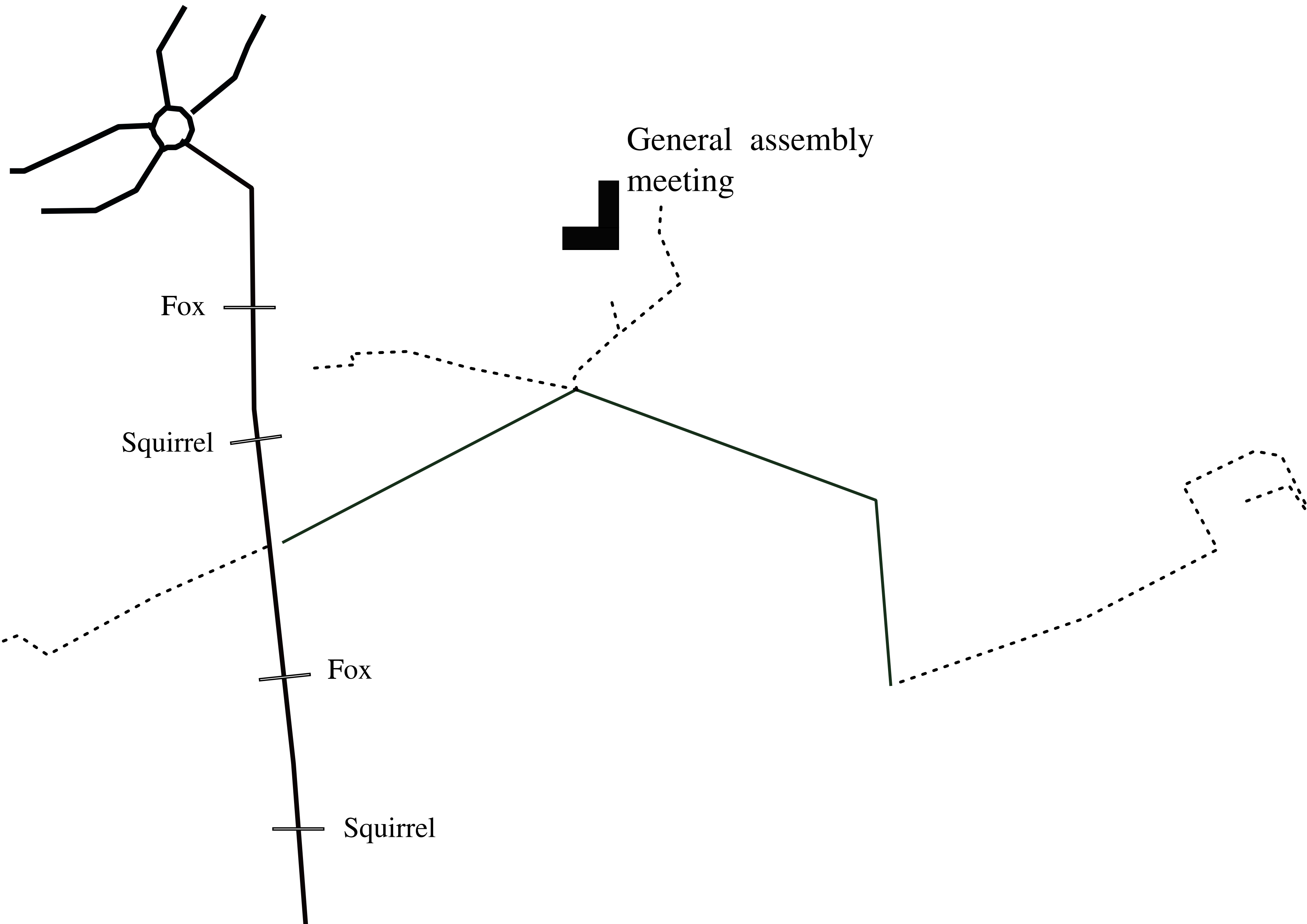
Figure A3: Map of the uninformed experiment (June session). Red fox Vulpes vulpes and red squirrel Sciurus vulgaris taxidermies were placed on each possible itinerary to the meeting location for the general assembly of the Bird Protection League (dashed roads are not accessible by car).
